# Skeletal dysplasia-causing TRPV4 mutations suppress the hypertrophic differentiation of human iPSC-derived chondrocytes

**DOI:** 10.1101/2021.06.15.448562

**Authors:** Amanda R. Dicks, Grigory I. Maksaev, Zainab Harissa, Alireza Savadipour, Ruhang Tang, Nancy Steward, Wolfgang Liedtke, Colin G. Nichols, Chia-Lung Wu, Farshid Guilak

## Abstract

Mutations in the TRPV4 ion channel can lead to a range of skeletal dysplasias. However, the mechanisms by which TRPV4 mutations lead to distinct disease severity remain unknown. Here, we use CRISPR-Cas9-edited human induced pluripotent stem cells (hiPSCs) harboring either the mild V620I or lethal T89I mutations to elucidate the differential effects on channel function and chondrogenic differentiation. We found that hiPSC-derived chondrocytes with the V620I mutation exhibited increased basal currents through TRPV4. However, both mutations showed more rapid calcium signaling with a reduced overall magnitude in response to TRPV4 agonist GSK1016790A compared to wildtype. There were no differences in overall cartilaginous matrix production, but the V620I mutation resulted in reduced mechanical properties of cartilage matrix later in chondrogenesis. mRNA sequencing revealed that both mutations upregulated several anterior *HOX* genes and downregulated antioxidant genes *CAT* and *GSTA1* throughout chondrogenesis. BMP4 treatment upregulated several essential hypertrophic genes in WT chondrocytes; however, this hypertrophic maturation response was inhibited in mutant chondrocytes. These results indicate that the TRPV4 mutations alter BMP signaling in chondrocytes and prevent proper chondrocyte hypertrophy, as a potential mechanism for dysfunctional skeletal development. Our findings provide potential therapeutic targets for developing treatments for TRPV4-mediated skeletal dysplasias.

## INTRODUCTION

Skeletal dysplasias comprise a heterogeneous group of over 450 bone and cartilage diseases with an overall birth incidence of 1 in 5000 (Krakow & Rimoin, 2010; Nemec et al., 2012; Ngo, Thapa, Otjen, & Kamps, 2018; Orioli, Castilla, & Barbosa-Neto, 1986; Superti-Furga & Unger, 2007). In the specific cases of moderate autosomal-dominant brachyolmia and severe metatropic dysplasia, among other dysplasias, arthropathies, and neuropathies, the disease is caused by mutations in transient receptor potential vanilloid 4 (TRPV4), a non-selective cation channel (Andreucci et al., 2011; Sun, 2012). For example, a V620I substitution (exon 12, G858A) in TRPV4 is responsible for moderate brachyolmia, which exhibits short stature, scoliosis, and delayed development of deformed bones (Kang, Shin, Auh, & Chun, 2012; Rock et al., 2008; Sun, 2012). These features, albeit more severe, are also present in metatropic dysplasia. Metatropic dysplasia can be caused by a TRPV4 T89I substitution (exon 2, C366T) and leads to joint contractures, disproportionate measurements, and, in severe cases, death due to small chest size and cardiopulmonary compromise (Camacho et al., 2010; Kang et al., 2012; Sun, 2012). Both V620I and T89I TRPV4 mutations are considered gain-of-function variants (Leddy, McNulty, Lee, et al., 2014; Loukin, Su, & Kung, 2011). Given the essential role of TRPV4 during chondrogenesis (Muramatsu et al., 2007) and cartilage homeostasis (O’Conor, Leddy, Benefield, Liedtke, & Guilak, 2014), it is hypothesized that TRPV4 mutations may affect the cartilaginous phase of endochondral ossification during skeletal development.

Endochondral ossification is a process by which bone tissue is created from a cartilage template (Breeland, Sinkler, & Menezes, 2021; Camacho et al., 2010; Krakow & Rimoin, 2010; Rimoin et al., 2007). During this process, chondrocytes transition from maintaining the homeostasis of cartilage, regulated by transcription factor SRY-box containing gene 9 (*SOX9*) (Breeland et al., 2021; Nishimura, Hata, Ono, et al., 2012; Prein & Beier, 2019; Sophia Fox, Bedi, & Rodeo, 2009), to hypertrophy. Hypertrophy is driven by runt related transcription factor 2 (*RUNX2*) and bone morphogenic protein (BMP) signaling (Breeland et al., 2021; Nishimura, Hata, Ono, et al., 2012; Prein & Beier, 2019) and leads to chondrocyte apoptosis or differentiation into osteoblasts to form bone (Breeland et al., 2021; Nishimura, Hata, Ono, et al., 2012; Prein & Beier, 2019). However, how TRPV4 and its signaling cascades regulate endochondral ossification remains to be determined.

The activation of TRPV4 increases *SOX9* expression (Muramatsu et al., 2007) and prevents chondrocyte hypertrophy and endochondral ossification (Amano et al., 2009; Hattori et al., 2010; Lui et al., 2019; Nishimura, Hata, Matsubara, Wakabayashi, & Yoneda, 2012; Nishimura, Hata, Ono, et al., 2012). One study found that overexpressing wildtype *Trpv4* in mouse embryos increased intracellular calcium (Ca^2+^) concentration and delayed bone mineralization (Weinstein, Tompson, Chen, Lee, & Cohn, 2014), a potential link between intracellular Ca^2+^, such as with gain-of-function TRPV4 mutations, and delayed endochondral ossification. Our previous study also observed increased expression of follistatin (*FST),* a potent BMP inhibitor, and delayed hypertrophy in porcine chondrocytes overexpressing human V620I- and T89I-TRPV4 (Leddy, McNulty, Guilak, & Liedtke, 2014; Leddy, McNulty, Lee, et al., 2014). While previous studies have greatly increased our knowledge of the influence of TRPV4 mutations on chondrogenesis and hypertrophy, most of them often involved animal models (Leddy, McNulty, Lee, et al., 2014; Weinstein et al., 2014) or cells (Camacho et al., 2010; Krakow & Rimoin, 2010; Loukin et al., 2011; Rock et al., 2008) (Leddy, McNulty, Lee, et al., 2014) overexpressing mutant TRPV4. Therefore, these approaches may not completely recapitulate the effect of TRPV4 mutations on human chondrogenesis.

Human induced pluripotent stem cells (hiPSCs), which are derived from adult somatic cells (Takahashi et al., 2007), offer a system for modeling human disease to study the effect of mutations throughout differentiation (Shaunak S. Adkar et al., 2017; Lee et al., 2021). In fact, two studies have used patient-derived hiPSCs with TRPV4 mutations to study lethal and non-lethal metatropic dysplasia-causing variants I604M (Saitta et al., 2014) and L619F (Nonaka et al., 2019), respectively. However, patient samples are often challenging to procure due to the rarity of skeletal dysplasias. In this regard, CRISPR-Cas9 technology allows the creation of hiPSC lines harboring various mutations along with isogenic controls (i.e., wildtype; WT).

The goal of this study was to elucidate the detailed molecular mechanisms underlying how two TRPV4 gain-of-function mutations lead to strikingly distinct severities of skeletal dysplasias (i.e., moderate brachyolmia vs. lethal metatropic dysplasia). To achieve this goal, we used CRISPR-Cas9 gene-edited hiPSC lines bearing either the V620I or T89I TRPV4 mutation, and their isogenic WT control, to delineate the effects of TRPV4 mutations on chondrogenesis and hypertrophy using RNA sequencing and transcriptomic analysis. We further examined the effects of the mutations on channel function and matrix production and properties. We hypothesized the V620I and T89I TRPV4 mutations would enhance chondrogenesis with differing degrees of altered hypertrophy. This study will improve our understanding of the role of TRPV4 in chondrocyte homeostasis and maturation and lay the foundation for treatment and prevention of TRPV4-mediated dysplasias.

## RESULTS

### Mutant TRPV4 has altered response to chemical agonist GSK101

We first assessed TRPV4 channel function and alterations in Ca^2+^ signaling due to the V620I and T89I mutations in day-28 hiPSC-derived chondrocytes using electrophysiology and fluorescence imaging. Using whole-cell patch clamping, we measured the basal membrane current of the hiPSC-derived chondrocytes from the mutated and WT lines. V620I-TRPV4 had the highest basal currents at both 70 and -70 mV (70/-70 mV pA/pF – WT: 18.52/5.93 vs. V602I: 77.79/55.33 vs. T89I: 40.97/50.13; Fig. 1A). However, when TRPV4 was inhibited with GSK205 (Kanju et al., 2016), TRPV4-specfic chemical antagonist, the three lines had similar, decreased currents (70/-70 mV – WT: 18.72/14.36 pA/pF vs. V620I: 13.55/9.15 pA/pF vs. T89I: 29.27/13.8 pA/pF; Fig. 1A). To capture the specific current through TRPV4, we took the difference of the basal current (no GSK205) and the average TRPV-inhibited current (with GSK205). TRPV4 inhibition caused a significant change in current in V620I at both 70 and -70 mV (70 mV – V620I: Δ64.28 vs. WT: Δ -0.19, p=0.0379 and T89I: Δ11.67, p<0.0001; -70 mV – V620I: Δ46.13 vs. WT: Δ -8.47, p<0.0001 and T89I: Δ36.33, p=0.0057; Fig. 1B). Interestingly, T89I-TRPV4 was not significantly different from WT despite also causing a gain-of-function in recombinant channels (Loukin et al., 2011).

**Figure 1.**
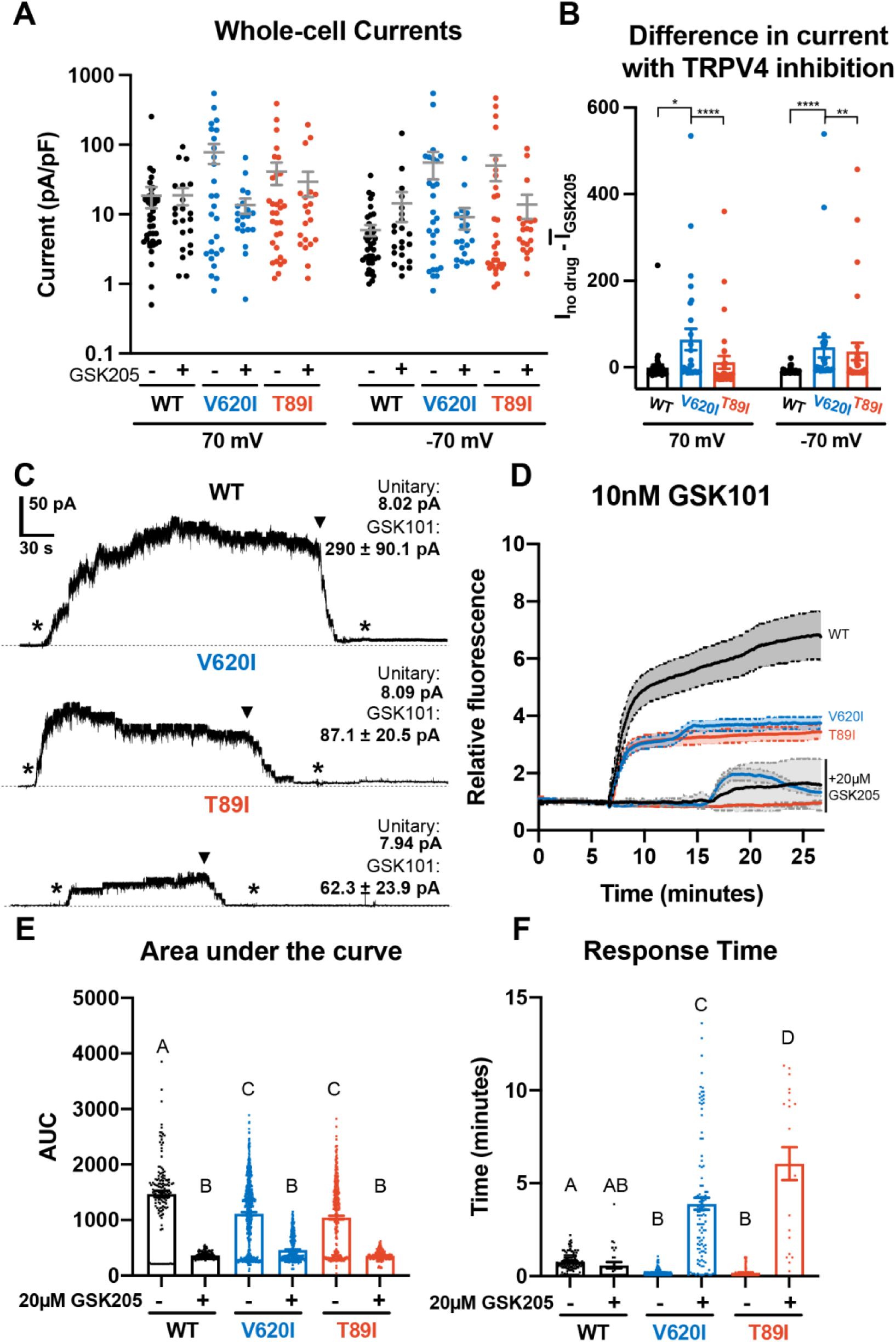
Differences in TRPV4 electrophysiological properties of WT and mutant hiPSC-derived chondrocytes. **A.** Whole-cell currents were higher, on average, in mutant hiPSC-derived chondrocytes than WT at 70 and -70 mV. TRPV4 inhibition with 20 µM GSK205 reduced mutant currents to similar levels as WT. Mean ± SEM. n=20-40 cells from 4 differentiations. **B.** The difference between the current through TRPV4 without GSK205 from the average current through inhibited channels was significantly higher in V620I. There was no difference between no drugs and GSK205 in WT. Mean ± SEM. n=27-40 from 4 differentiations. Kruskal-Wallis test with multiple comparisons comparing cell lines at 70 mV and -70 mV. *p<0.05, **p<0.01, ****p<0.001. **C.** Inside-out excised patches of WT had a higher current in response to 10 nM GSK101 (indicated by *) than mutants. The addition of 10 nM GSK101 + 20 µM GSK205 (indicated by ▾) decreased the current and continued to block the channel when GSK101 alone was re-introduced (*). Representative plots with average unitary current and current in response to GSK101. Mean ± SEM. N = 5, 9, and 8 for WT, V620I, and T89I, respectively from 2 differentiations. **D.** Mutant TRPV4 decreased the channels’ sensitivity to activation with GSK101 as shown with confocal imaging of ratiometric fluorescence indicating Ca^2+^ signaling. GSK205 attenuated GSK101-mediated signaling. Mean ± 95% CI. n = 3 experiments with a total of 158-819 cells per line. **E.** Quantification of the area under the curve of (D). Mean ± SEM. n = 158-819 cells from 3 experiments. Ordinary two-way ANOVA with Tukey’s post-hoc test. Interaction, cell line, and treatment p<0.0001. **F.** Time of initial response of each responding cell (≥25% of frames for that cell are responding) measured from the addition of stimulus. Mutant TRPV4 responded faster to GSK101, but the response was significantly slowed by GSK205. Responding frames were considered to have a fluorescence greater than the mean plus three times the standard deviation. Mean ± SEM. n=21-360 responding cells from 3 experiments. Ordinary two-way ANOVA with Tukey’s post-hoc test. Interaction, cell line, and treatment p<0.0001.

Next, we activated WT and mutant TRPV4 with chemical agonist GSK1016790A (GSK101) (Jin et al., 2011) and found that the mutations decreased the cellular response to the agonist, resulting in reduced Ca^2+^ signaling. These results were supported using two methods: inside-out excised patches and confocal imaging of Ca^2+^ signaling (Fig. 1C-D). The representative traces of inside-out patches showed increased current through the patch with the addition of GSK101 and the attenuation by GSK205 (Fig. 1C). GSK205 continued to block the channel and prevented another increase in current despite the addition of GSK101. Though the unitary currents were indistinguishable (8 pA at -30mV) among WT and mutants, in excised inside-out patches WT typically produced higher GSK101-induced currents than the mutants (WT: 290 pA vs. V620I: 87.1 pA and T89I: 62.3 pA at -30mV), potentially indicative of more channels per patch (Fig. 1C). In the confocal imaging experiments, a ratiometric fluorescence indicated Ca^2+^ signaling of the hiPSC-derived chondrocytes in response to either 10 nM GSK101 or a cocktail of 10 nM GSK101 and 20 µM GSK205. WT cells had significantly higher fluorescence, and therefore Ca^2+^ signaling, in response to GSK101 according to the plots and their area under the curve (AUC; WT: 1470 vs. V620I: 1114 and T89I: 1044; p<0.0001; Fig. 1D-E). The presence of GSK205 attenuated this response for all three lines, confirming the Ca^2+^ influx was due to the TRPV4 ion channel (WT: 366 vs. V620I: 460 vs T89I: 358). We also evaluated the response time of the cells to GSK101 and GSK101+GSK205. We considered a cell to be responding if more than a quarter of its frames, after stimuli, had a fluorescence higher than the mean baseline plus 3 times the standard deviation. The mutants responded faster to GSK101 than the WT (WT: 46.2 sec vs. V620I: 12 sec, p=0.0048 and T89: 10.8 sec, p=0.0097; Fig. 1F). Interestingly, the addition of GSK205 did not significantly slow the response of WT, but it did slow the response of the mutants, with the severe mutation slower than the moderate (WT: 35.4 sec vs. V620I: 234 sec and T89: 366 sec; p<0.0001; Fig. 1F).

### Chondrogenic differentiation of WT and mutant hiPSC lines

To confirm if the hiPSCs with dysplasia-causing mutations would undergo proper chondrogenesis, we differentiated CRISPR-Cas9-edited hiPSCs with mutant *TRPV4* alongside an isogenic wildtype (WT) using our previously published protocol (S. S. Adkar et al., 2019; Wu et al., 2021). After 12 days of monolayer mesodermal differentiation, the cells underwent 42 days of chondrogenic differentiation, and pellets were collected at days 7, 14, 28, and 42. At day 28, the three lines had similar chondrogenic matrix as shown with Safranin-O staining for sulfated glycosaminoglycans (sGAGs) and collagen type 2 alpha chain 1 (COL2A1) labeling with immunohistochemistry (IHC; Fig. 2A-B). All three lines had little to no labeling of fibrocartilage marker COL1A1 and hypertrophic cartilage marker COL10A1 with IHC (Fig. 2C-D). To quantitatively confirm the matrix production throughout chondrogenesis, we performed biochemical assays to measure sGAG production and normalize it to double-stranded DNA content. As expected, differences in matrix production were significant between time points (p<0.0001; Fig. 2E). The sGAG/DNA ratio increased in WT by 8-fold and in V620I and T89I by 5- to 5.5-fold from day 14 to 28 (p<0.0001). V620I pellets also increased in matrix content by 150% from day 28 to 42 (p=0.0163) with all three lines reaching an sGAG/DNA ratio of approximately 30. However, there were no differences in sGAG/DNA ratios among the three cell lines at any time point (cell line: p=0.1206; interaction: p=0.7426).

**Figure 2.**
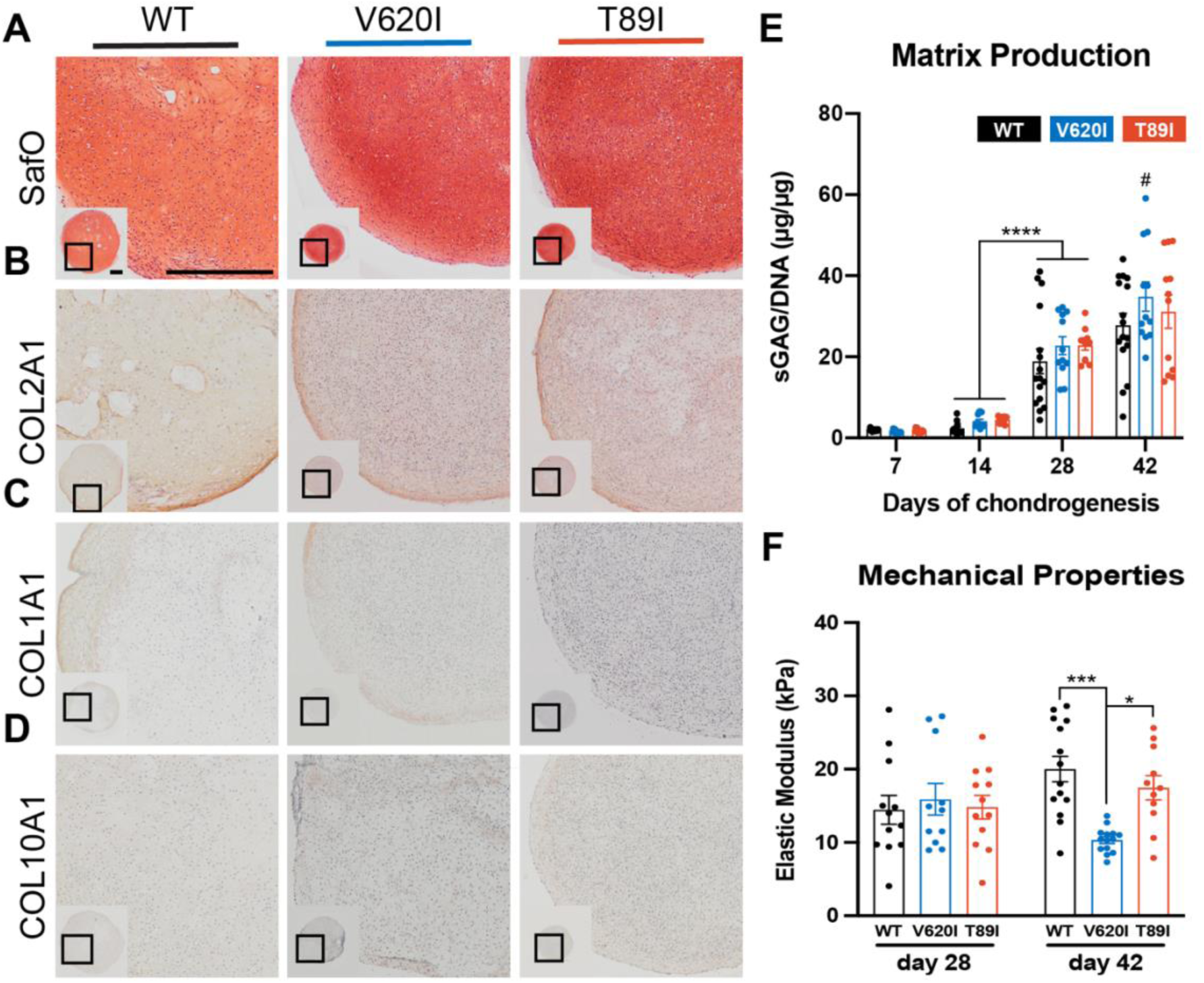
Mutant TRPV4 had little effect on chondrogenic matrix production. **A-D**. WT, V620I, and T89I day-28 pellets exhibit similar matrix production shown by staining for sGAGs with Safranin-O and hematoxylin (**A**) and labeling with IHC for COL2A1 (**B**), COL1A1 (**C**), and COL10A1 (**D**). Scale bar = 500 µm. Representative images from 3-4 differentiations. **E**. The sGAG/DNA ratio increased in all three lines from day 14 to 28 of chondrogenesis. There were no differences between lines at each time point. Mean ± SEM. n = 11-16 from 3-4 differentiations. ****p<0.0001 Statistical significance determined by an ordinary two-way ANOVA with Tukey’s post-hoc test. **F.** There were no differences in the elastic modulus of the matrix at day 28. Day-42 V620I had a significantly lower elastic modulus than WT and T89I. Mean ± SEM. n=11-14 from 3 experiments. *p<0.05, ***p<0.001 Statistical significance determined by an ordinary two-way ANOVA with Tukey’s post-hoc test.

Atomic force microscopy (AFM) was then used to measure the mechanical properties of the hiPSC-derived cartilaginous matrix deposited by the WT and two TRPV4 mutated cell lines. The elastic modulus ranged from 14 to 20 kPa, consistent with mouse iPSC-derived cartilage (Diekman et al., 2012). At day 28, the three lines had similar properties (WT: 14.4 kPa vs. V620I: 15.9 kPa vs. T89I: 14.8 kPa); however, at day 42, V620I had a significantly decreased elastic modulus (V620I: 10.32 kPa vs. WT: 20.0 kPa, p=0.0004 and T89I: 17.5 kPa, p=0.0328; Fig. 2F). These experiments indicated that all three lines could properly differentiate into chondrocytes and had similar cartilaginous matrix production by day 28. Therefore, we used the day 28 time point for further studies.

### TRPV4 mutations altered chondrogenic gene expression in hiPSC-derived chondrocytes

RT-qPCR analysis throughout differentiation shows that mutants had higher *ACAN* expression compared to WT at day 28; however, expression decreased at day 42 in T89I (day-42 fold changes; V620I: 5933 vs. WT: 2687, p=0.0016 and T89I: 2631, p=0.0058; Fig. 3A). *COL2A1* expression was similar among the three lines at day 28 but significantly lower in T89I at day 42 (day-42 fold changes; T89I: 2798 vs. WT: 9209, p=0.0144 and V620I: 7177, p=0.0007; Fig. 3B). Throughout chondrogenesis, V620I significantly increased expression of chondrogenic transcription factor *SOX9* (day-42 fold changes; V620I: 195.3 vs. WT: 44.29, p<0.0001 and T89I: 32.19, p=0.0003; Fig. 3C) and *TRPV4* (day-42 fold changes; V620I: 168.5 vs. WT: 48.82, p<0.0001 and T89I: 44.72, p<0.0001; Fig. 3D). On the other hand, T89I significantly increased expression of pro-inflammatory, calcium binding protein *S100B* (Yammani, 2012) throughout chondrogenesis (day-42 fold changes; T89I: 1363 vs. WT: 362.0, p=0.0018 and V620I: 507.8, p=0.0439; Fig. 3E). T89I also had significantly higher expression of fibrocartilage marker *COL1A1* at days 7, 14, and 28 than the other two lines, and both mutations had increased expression at day 42 compared to WT (day-42 fold changes; WT: 32.47 vs. V620I: 76.42, p=0.01.58 and T89I: 74.23, p=0.0132; Fig. 3F). In contrast, hypertrophic marker *COL10A1* was significantly higher in the WT line than the mutants at days 28 and 42 (day-42 fold changes; WT: 615.7 vs. V620I: 71.00, p=0.0001 and T89I: 83.07, p=0.0015; Fig. 3G). Surprisingly, there was not a significant increase in follistatin (*FST*) expression in mutants at later time points (day- 42 fold changes; WT: 0.5342 vs. V620I: 0.6808, p=0.6882 and T89I: 0.3158, p>0.9999; Fig. 3H).

**Figure 3.**
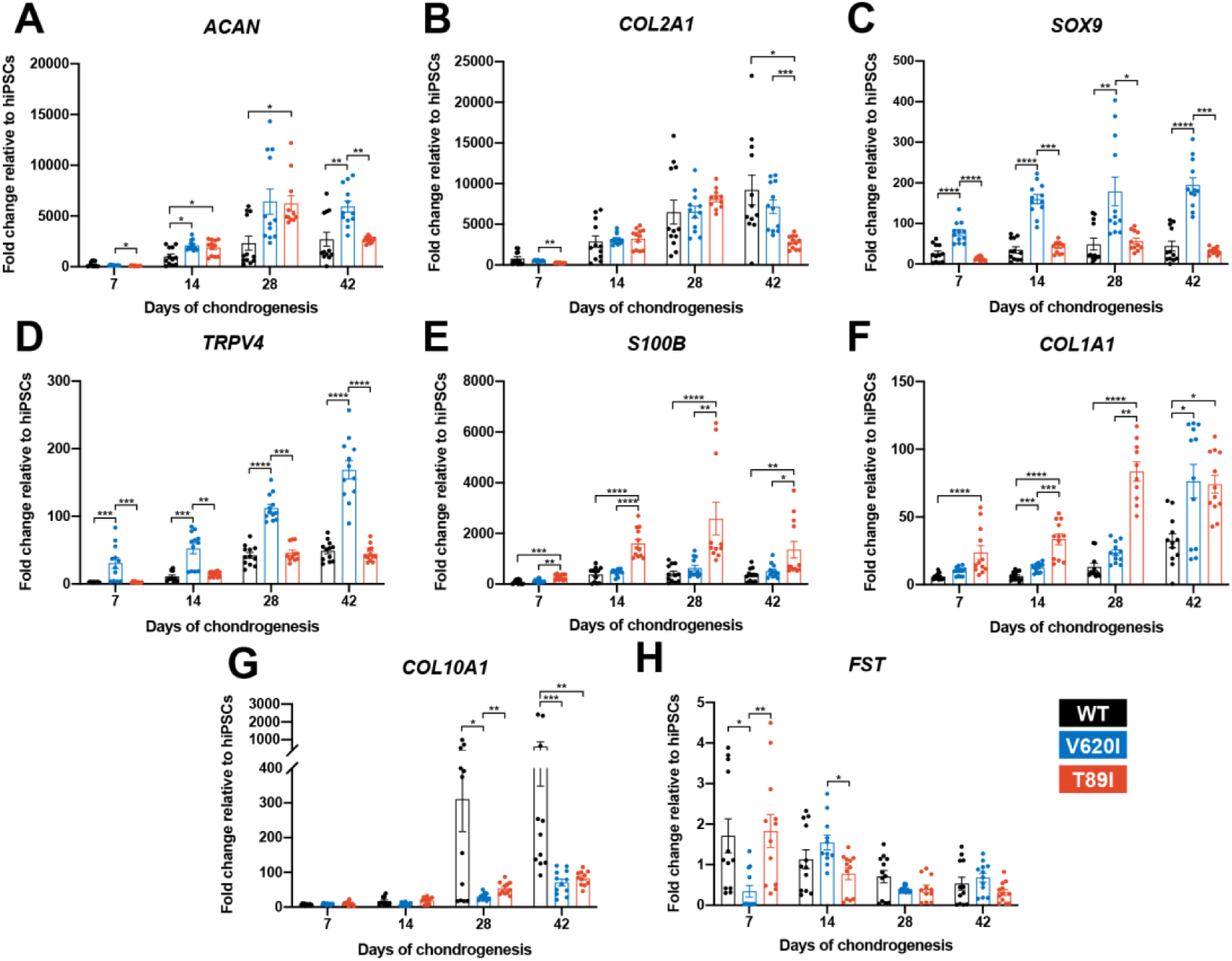
V620I and T89I had differing effects on gene expression during chondrogenic differentiation. **A.** T89I and V620I had increased *ACAN* gene expression at day 28 and 42, respectively, compared to WT. **B.** Day-42 T89I chondrocytes had decreased expression of *COL2A1*. **C-D.** V620I increased expression of *SOX9* (C) and *TRPV4* (D) throughout chondrogenesis. **E-F.** T89I increased expression of *S100B* (E) and *COL1A1* (F) throughout chondrogenesis. **G.** Both mutations decreased *COL10A1* gene expression at day 28 and 42, compared to WT. **H.** There were no differences in *FST* expression at later time points day 28 and 42. Mean ± SEM. n=10-12 from 3 differentiations. *p<0.05, **p<0.01, ***p<0.001, ****p<0.0001 Significance determined by one-way ANOVA with Tukey’s post-hoc test for each time point.

To obtain comprehensive transcriptomic profiles of WT and TRPV4 mutated cell lines, we performed bulk RNA sequencing of day-28 chondrogenic pellets. We compared V620I and T89I gene expression to WT and plotted the log_2_ fold change in heatmaps (Fig. 4A-B). While many chondrogenic and hypertrophic genes had similar levels of expression between the lines, the mutants had increased expression of cartilage ECM genes cartilage oligomeric matrix protein (*COMP*), collage type 6 alpha chains 1 and 3 (*COL6A1, COL6A3*), growth differentiation factor 5 (*GDF5),* high-temperature requirement A serine peptidase 1 *(HTRA1),* and secreted protein acidic and cysteine rich (*SPARC)* (Fig. 4A). In contrast, the mutants had decreased expression of hypertrophic markers *COL10A1,* secreted phosphoprotein 1 *(SPP1),* and alkaline phosphatase, biomineralization associated (*ALPL)* in addition to *SST* (Fig. 4B). The mutations up-regulated expression levels of bone morphogenic protein 6 (*BMP6*), transforming growth factor 3 (*TGFB3)*, nuclear factor of activated T-Cells C2 (*NFATC2*), Twist family BHLH transcription factor 1 (*TWIST1*), ADAM metallopeptidase with thrombospondin type 1 motif 4 (*ADAMTS4*), and *WNT3A* (Fig. 4B). The mutations also downregulated osteoblastogenesis transcription factors *SOX2* and *SOX11* and previously identified genes governing off-target differentiation during hiPSC chondrogenesis including nestin (*NES),* orthodenticle homeobox 2 (*OTX2*)*, WNT7A,* and *WNT7B* (Fig. 4B). Interestingly, *BMP4* was downregulated to a greater extent in V620I than T89I when compared to WT.

**Figure 4.**
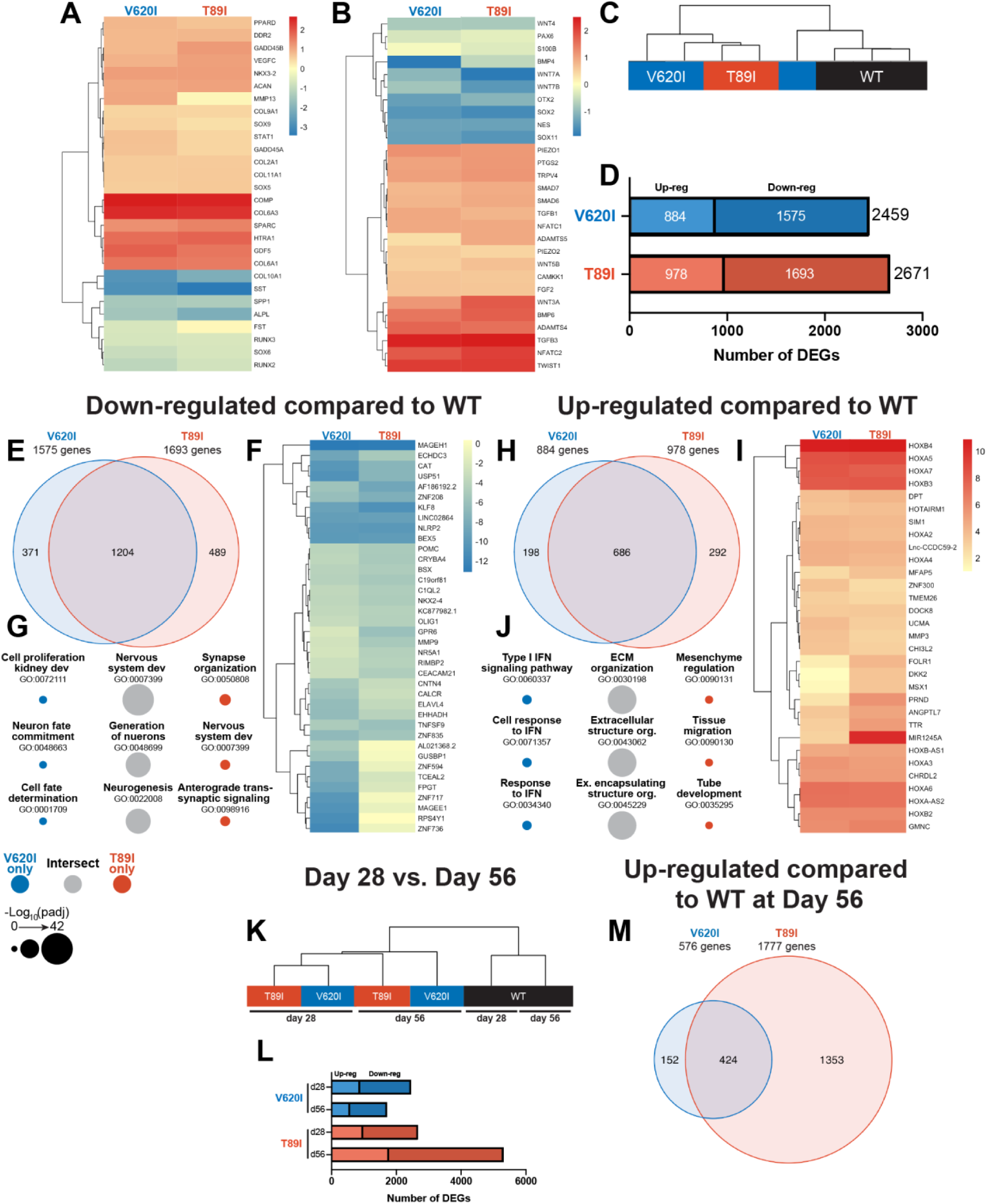
Dynamic changes in transcriptomic profiles of V620I and T89I mutants during chondrogenesis. **A-B.** Heatmaps comparing the log_2_ fold change of common chondrogenic and hypertrophic genes (A) and growth factor and signaling genes (B) in day-28 V620I and T89I chondrocytes compared to WT. **C.** Clustering of the samples using Euclidean distances reveals that V620I and T89I hiPSC-derived chondrocytes are more similar to each other than WT. **D.** The number of up- and down-regulated DEGs in V620I and T89I day-28 chondrocytes compared to WT. **E-G.** Analysis of the down-regulated genes compared to WT. **E.** A Venn diagram reveals the number of similar and different down-regulated DEGs between V620I and T89I, where most genes are shared. **F.** A heatmap showing the log_2_ fold change, compared to WT, of the top 25 down-regulated genes for each line. **G.** The top 3 GO Terms (biological process) associated with the DEGs unique to V620I, shared between V620I and T89I, and unique to T89I. **H-J.** Analysis of the up-regulated genes compared to WT. **H.** A Venn diagram reveals the number of similar and different up-regulated DEGs between V620I and T89I, where most genes are shared. **I.** A heatmap showing the log_2_ fold change, compared to WT, of the top 25 up-regulated genes for each line. **J.** The top 3 GO Terms (biological process) associated with the DEGs unique to V620I, shared between V620I and T89I, and unique to T89I. **K.** Clustering of the day-28 and day-56 samples using Euclidean distances reveals that the WT chondrocytes, at both day 28 and 56, cluster together while mutants cluster by time point. **L.** The number of up-regulated and down-regulated DEGs for V620I and T89I compared to WT at day 28 and day 56. **M.** A Venn diagram reveals the number of similar and different up-regulated DEGs between V620I and T89I, with T89I becoming more unique at day 56. n=3-4 samples.

### V620I and T89I mutants demonstrate similar gene expression profiles

First, to evaluate the similarities and differences in transcriptomic profiles between the hiPSC-derived chondrocytes with and without the TRPV4 mutations, we computed the Euclidean distance between day-28 samples of each cell line. The WT samples clustered away from the mutants, and the V620I samples were the most variable. (Fig. 4C). In terms of total differentially expressed genes (DEGs) compared to WT, V620I had 8% fewer DEGs than T89I (2459 vs. 2671; Fig. 4D). Mutants had only about half of the number of up-regulated genes compared to down-regulated genes (V620I: 884 vs. 1575, T89I: 978 vs. 1693; Fig. 4D). The majority of the down-regulated DEGs were shared between the two mutants when compared to WT, comprising 76% and 71% of V620I’s and T89I’s total down-regulated DEGs, respectively (Fig. 4E). We plotted the top 25 most down-regulated DEGs for each line in a heatmap. These included antioxidant catalase (*CAT*), anti-inflammatory nucleotide-binding and leucine-rich repeat receptor family pyrin domain containing 2 (*NLRP2),* and kruppel like factor 8 *(KLF8)* (Fig. 4F). Interestingly, many of the down-regulated DEGs, both unique and shared between V620I and T89I, were associated with Gene Ontology (GO) terms related to nervous system development, including many potassium channel genes (i.e., *KCN* family; Fig. 4G).

In contrast, 686 up-regulated DEGs were shared by both mutants, while 22% of V620I’s and 30% of T89I’s up-regulated DEGs were unique to each mutation (198 vs. 292; Fig. 4H). A heatmap of the top 25 up-regulated DEGs showed that several homeobox (HOX) genes were highly expressed in chondrocytes with the TRPV4 mutations (Fig. 4I). These included *HOXA2* to *HOXA7, HOXA-AS2, HOXB2* to *HOXB4,* and *HOXB-AS1,* which are associated with morphogenesis and anterior patterning (Seifert, 2015). Furthermore, the shared, up-regulated DEGs between two mutants are associated with extracellular matrix production and organization and growth factor binding in GO term analysis, while V620I genes were associated with type I interferon (Fig. 4J). These data highlight an early morphogenic genetic profile in hiPSC-derived chondrocytes with the V620I and T89I mutations.

### The severe T89I mutation inhibits chondrocyte hypertrophy more than moderate V620I mutation

Following an additional 4 weeks of chondrogenic culture, we performed RNA sequencing to investigate how the differences between the WT and the two mutants change with further differentiation. Using Euclidean distances, we compared the WT, V620I, and T89I hiPSC-derived chondrocytes at both day 28 and 56 (Fig. 4K). WT clustered together at both day 28 and 56; however, the mutants clustered by time point. Again, there were more down-regulated genes than up-regulated at day 56 (Fig. 4L). T89I had the most DEGs, and the number increased from day 28 to 56. In contrast, V620I’s DEGs decreased at day 56. 74% of V620I up-regulated DEGs, but only 24% of T89I DEGs, were shared between the two lines (424 total genes; Fig. 4M). These intersecting, up-regulated genes were associated with the biological processes of skeletal development, morphogenesis, and patterning due to the up-regulation of many *HOX* genes (Fig. 4 – Fig. S1A). Most of the top up- and down-regulated genes were consistent between day 28 and 56 (Fig. 4 – Fig. S1A-B), including both anterior and posterior *HOX* genes (i.e., *HOXA1* to *HOXA7, HOXB2* to *HOXB4, HOXB6* to *HOXB8, HOXC4, HOXD8, HOXA-AS2-3,* and *HOXB-AS1-2*)(Seifert, 2015). Although V620I and T89I TRPV4 mutants continued to share the up-regulated *HOX* genes, which may be responsible for dysfunctional chondrogenic hypertrophy compared to WT cells, our results also indicate that these two mutated lines started to demonstrate divergent transcriptomic profiles in later chondrogenesis.

### TRPV4 mutations exhibit dysregulated BMP4-induced chondrocyte hypertrophy

To evaluate how TRPV4 mutations may affect hypertrophy, BMP4 was added to the chondrogenic medium with and without TGFβ3 to stimulate hypertrophic differentiation starting at day 28 of chondrogenic pellet culture (Craft et al., 2015). At day 56, Safranin-O staining indicated the BMP4-treated WT had developed a more hypertrophic phenotype compared to TGFβ3- and TGFβ3+BMP4-treated pellets with enlarged chondrocytes (cell diameter; WT-BMP4: 27.6 µm vs. WT-TGFβ3: 11.8 µm, V620I-BMP4: 12.5 µm, and T89I-BMP4: 11.3 µm; p<0.0001; Fig. 5A-B). This phenotype was not present in any of the groups from the V620I and T89I lines. Western blot further confirmed this with an increase in hypertrophic cartilage protein COL10A1 secretion in the WT-BMP4 group (Fig. 5C), consistent with the day-28 gene expression data (Fig 3G, 4A). RNA sequencing and PCA revealed that the WT line was more sensitive to BMP4, as indicated by the arrows (Fig. 5D). Given that the BMP4-treated WT chondrocytes had the most apparent hypertrophic phenotype, later analyses were performed comparing the BMP4- and TGFβ3-treated chondrocytes for simplification.

**Figure 5.**
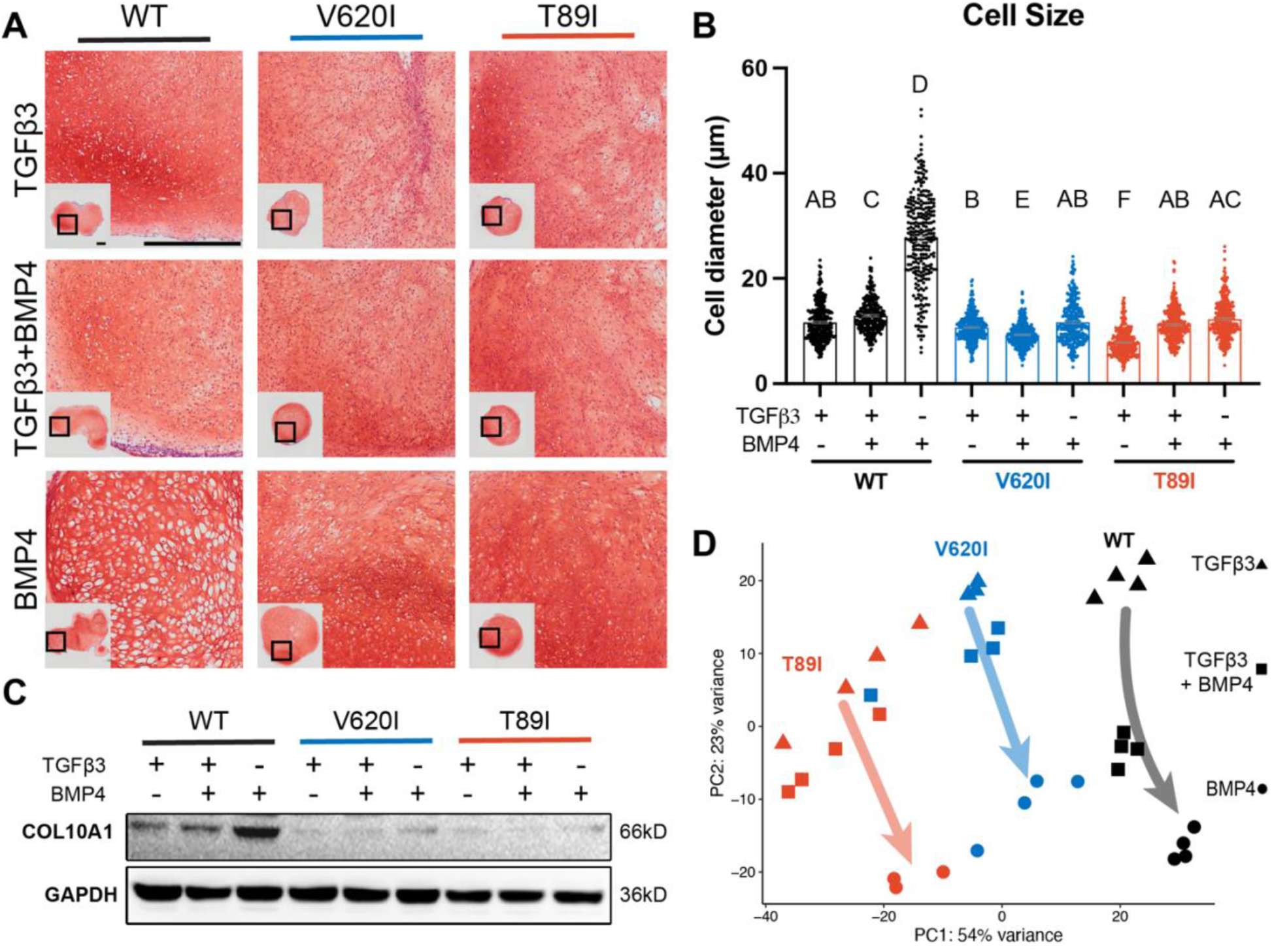
WT chondrocytes are more sensitive to BMP4 treatment. **A.** WT chondrocytes treated with BMP4 developed a hypertrophic phenotype with enlarged lacunae, which was not present in the mutant cell lines or other conditions, as shown by Safranin-O and hematoxylin staining. Scale bar = 500 µm. Representative images from 2 experiments. **B.** Cell diameter was significantly increased in the WT with BMP4 treatment compared to all other groups indicating a hypertrophic phenotype. Mean ± SEM. n = 249-304 cells from 2 pellets. Different letters indicate statistical significance (p<0.05) between groups as determined by Kruskal-Wallis test with multiple comparisons since data was not normally distributed. **C.** Western blot shows that WT had increased production COL10A1 in response to BMP4 treatment. **D.** PCA of bulk RNA-seq reveals an increased sensitivity to BMP4 (and TGFβ3+BMP4) treatment in WT hiPSC-derived chondrocytes compared to V620I and T89I. n=3-4 samples.

Hierarchical K-means clustering of gene expression profiles of BMP4- and TGFβ3- treated chondrocytes resulted in 9 unique clusters, as determined using the gap statistics method (Fig. 6A). Most of the clusters, including the largest (i.e., cluster 1), showed up-regulation of gene expression with BMP4 treatment, while clusters 4, 5, and 9 showed down regulation. The gene expression per group for each cluster is listed in Supplemental File 1. Overall, WT responded to BMP4 treatment with the largest number of DEGs, over 2500, with only 22% of them shared among all three lines (Fig. 6B). Although cluster 1 shows an overall increase in gene expression with BMP4 treatment, WT had a larger increase in expression than the mutants (Fig. 6C). In fact, some of the genes that were up-regulated with BMP4 treatment in WT may have no change or down-regulation in mutants (cluster 1, Fig. 6A).

**Figure 6.**
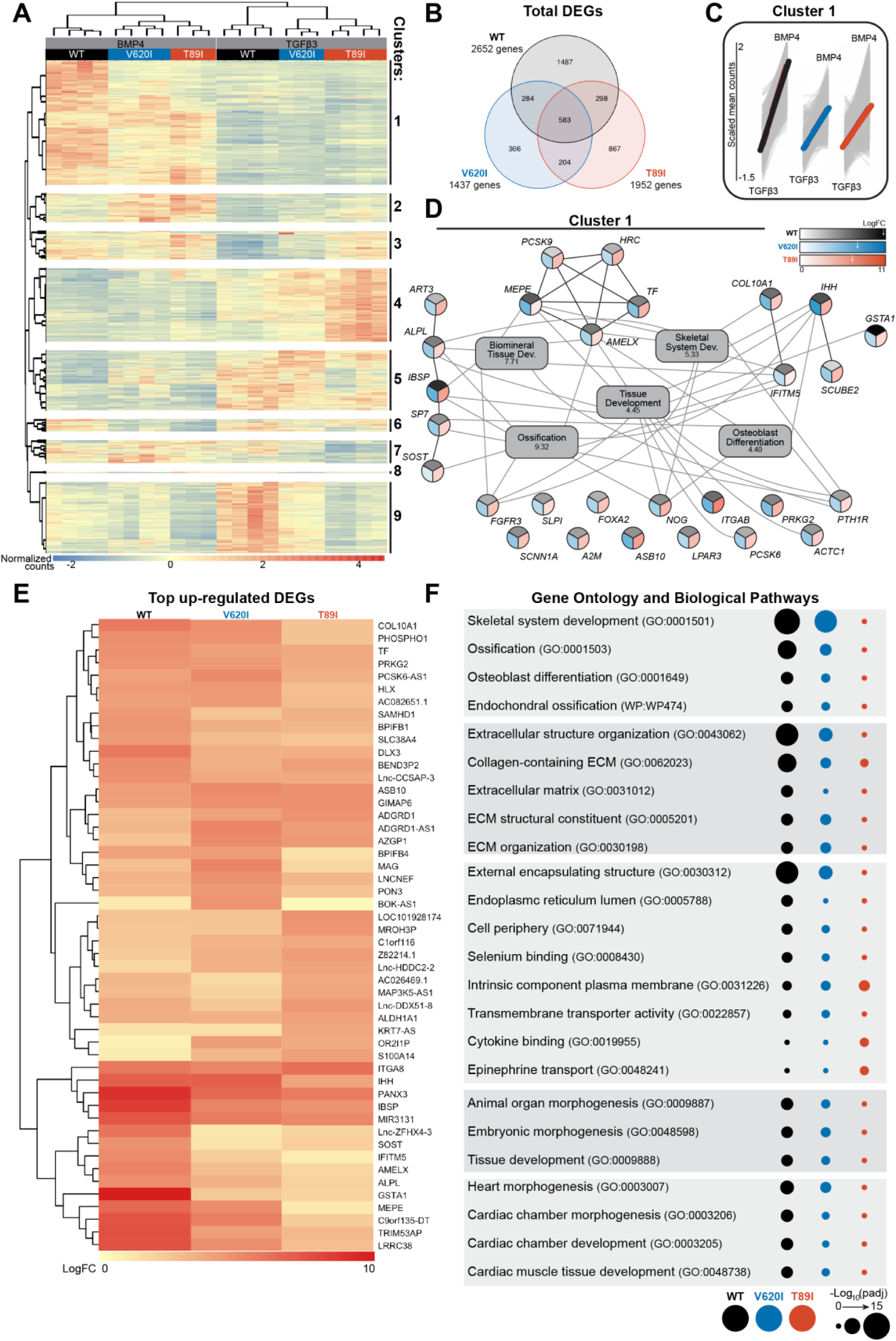
V620I and T89I had an inhibited hypertrophic response to BMP4 treatment. **A.** There are 9 clusters of genes based on expression and hierarchical k-means clustering of the samples. **B.** Venn diagram shows similar and distinct DEGs in response to BMP4 treatment in all three lines. **C.** Cluster 1 represented increasing in expression from TGFβ3-treatment to BMP4-treatment (left to right on x-axis). Y-axis scale (-1.5 to 2) represents the scaled mean counts. **D.** A protein-protein interaction network with functional enrichment analysis of cluster 1 reveals the top regulating genes and their associated concepts. Connections between protein-coding genes and GO processes are based on the average log fold change between cell lines. Coloring of the protein-coding gene circles is divided into three to represent the log fold change for each cell line as shown in the legend. The white arrows in the legend indicates the location of the maximum log fold change for each respective cell line. The grey boxes represent the top 5 GO terms (biological process) identified for the network with the log_10_(false discovery rate) underneath the term. **E.** A heatmap of the top 25 up-regulated genes, and their log_2_ fold change, in each line compared to their respective TGFβ3 controls. **F.** The top GO terms and biological pathways associated with the up-regulated DEGs with BMP4 treatment. Symbol color represents the cell line, and size represents the -log_10_(p_adj_).

As cluster 1 represents the primary response to BMP4 treatment and may highlight how the TRPV4 mutations inhibit chondrocyte hypertrophy, we constructed a gene network of this cluster (Fig. 6D). The log fold change of each gene per cell line is represented by a color scale, which is consistent with WT having overall higher expression of the genes (as indicated by the white arrows in the legend; Fig. 6D). With GO term analysis, the cluster 1 gene network is highly associated with ossification, biomineral tissue development, skeletal system development, tissue development, and osteoblast differentiation (Fig. 6D). Alkaline phosphatase, biomineralization associated (*ALPL*), amelogenin X-linked (*AMELX*), fibroblast growth factor receptor 3 (*FGFR3*), interferon induced transmembrane protein 5 (*IFITM5*), Indian hedgehog (*IHH*), parathyroid hormone 1 receptor (*PTH1R*), and noggin (*NOG*) were connected to at least 4 of the top 5 GO terms. Of those, *ALPL, AMELX,* and *IFITM5* showed much higher expression in WT than the mutants alongside antioxidant glutathione S-transferase alpha 1 (*GSTA1*) and bone ECM proteins integrin binding sialoprotein (*IBSP*) and matrix extracellular phosphoglycoprotein (*MEPE*). Lack of expression of these key genes may be responsible for the inhibited hypertrophy in TRPV4 V620I- and T89I-mutated chondrocytes.

We next investigated and plotted the top 25 up-regulated genes for each line with BMP4 treatment (compared to their respective TGFβ3 control) (Fig. 6E). 88% of these genes were also present in cluster 1. The key genes *ALPL, AMELX, IFITM5, GSTAI, IBSP*, and *MEPE* had distinctly higher expression in WT than mutants, in agreement with the network analysis. Both mutants showed higher expression than WT of ankyrin repeat and SOCS box containing 10 (*ASB10*), GTPase, IMAP family member 6 (*GIMAP6*), and adhesion G protein-coupled receptor D1 (*ADGRD1*) when compared to their corresponding TGFβ3 control group. GO term analysis was further performed on all BMP4 up-regulated DEGs for each line (Fig 6F). WT was highly associated with skeletal system development, ossification, endochondral ossification, and extracellular structure organization, followed by V620I mutants, while T89I showed little to no association with these concepts.

## DISCUSSION

To elucidate the detailed molecular mechanisms underlying the distinct severity of skeletal dysplasias caused by two TRPV4 mutations (brachyolmia-causing V620I vs. metatropic dysplasia-causing T89I), we used CRISPR-Cas9 gene editing to generate hiPSC-derived chondrocytes bearing these two mutations. We observed that day-28 chondrocytes exhibited differences in channel function and gene expression between the mutants and WT control. Differences in transcriptomic profiles between V620I and T89I and from WT became more apparent with maturation following 4 additional weeks of culture with TGFβ3 or hypertrophic differentiation with BMP4 treatment. Of note, WT was significantly more sensitive to BMP4- induced hypertrophy. At the transcriptomic level, TRPV4 mutations inhibited chondrocyte hypertrophy, particularly with the T89I mutation, whereas V620I exhibited a milder phenotype, consistent with the clinical presentation of these two conditions. Our results suggest that skeletal dysplasias may be, at least in part, resulting from improper chondrocyte hypertrophy downstream of altered TRPV4 function. Furthermore, with our genome-wide RNA sequencing analysis, we also identified several putative genes that may be responsible for these dysregulated pathways in human chondrocytes bearing V620I or T89I TRPV4 mutations.

Our findings are generally consistent with previous non-human models of V620I and T89I mutations. Two other models that have studied the V620I and T89I mutations include *X. laevis* oocytes injected with rat TRPV4 cRNA (Loukin et al., 2011) or primary porcine chondrocytes transfected with human mutant TRPV4 (Leddy, McNulty, Lee, et al., 2014). Both reports and our current study investigated the baseline currents of the mutant TRPV4 compared to WT. Here, we used patch clamping and observed high basal currents in V620I with a significant decrease when TRPV4 was inhibited. However, this characteristic was trending, but not significant, in T89I, despite both V620I and T89I being reported as gain-of-function mutations (Camacho et al., 2010; Rock et al., 2008). Both the *X. laevis* oocyte and porcine chondrocyte models confirmed high basal currents through V620I-TRPV4 (Leddy, McNulty, Lee, et al., 2014; Loukin et al., 2011). Interestingly, *X. laevis* oocytes, but not the humanized porcine chondrocytes, showed an increase in basal Ca^2+^ signaling through T89I (Leddy, McNulty, Lee, et al., 2014; Loukin et al., 2011). Furthermore, our results were consistent with a summary of TRPV4 channelopathies reporting an increase in conductivity in V620I but no change in T89I (Sun, 2012). The conflicting basal current results could be due to the species of the TRPV4, but this was not the case regarding channel activation. As mentioned, the hiPSC-derived chondrocytes with V620I and T89I TRPV4 had reduced currents and Ca^2+^ signaling in response to chemical agonist GSK101. However, our previous study showed the porcine chondrocytes with mutant human TRPV4 had increased peak Ca^2+^ signaling in response to hypotonic changes (Leddy, McNulty, Lee, et al., 2014). This discrepancy could be due to the mode of activation of TRPV4 (i.e., osmotic vs. chemical agonist). In contrast, the oocytes with mutant rat TRPV4 had lower currents in response to both hypotonic and chemical (GSK101) TRPV4 activation compared to WT-TRPV4, consistent with our findings. It can be speculated that there is decreased sensitivity to the antagonist because the mutated hiPSC-derived chondrocytes are compensating for the increased basal activity by reducing the number of TRPV4 channels, or other ion channels and signaling transducers as shown with the RNAseq data and associated GO terms. The increased basal currents and decreased channel sensitivity to TRPV4 agonist GSK101 with mutated TRPV4 is also likely due to an increased open probability of TRPV4 making the channels less likely to open due to an agonist (Loukin et al., 2011). The obvious differences in both resting and activated states confirm functional differences with TRPV4 mutations that may ultimately lead to changes downstream of the channel, which alter joint development and result in skeletal dysplasias.

It was hypothesized, in the porcine chondrocyte study, that the increased Ca^2+^ signaling due to the V620I and T89I TRPV4 mutations increased *FST* expression that inhibited BMP signaling and hypertrophy (Leddy, McNulty, Guilak, et al., 2014; Leddy, McNulty, Lee, et al., 2014). Surprisingly, we found no differences in *FST* expression in mutant hiPSC-derived chondrocytes compared to WT. However, our previous study used non-human cells, which could alter the effects of the human TRPV4 mutations and downstream gene expression. Another previous hypothesis made was that the altered TRPV4 signaling increased *SOX9* expression, a known regulator of resting and proliferating chondrocytes upregulated by TRPV4 activation (Muramatsu et al., 2007), thus decreasing hypertrophy (Rock et al., 2008). *SOX9*-knockin mice exhibit a dwarfism phenotype (Amano et al., 2009), and *SOX9* overexpression inhibits hypertrophy and endochondral ossification (Hattori et al., 2010; Lui et al., 2019), likely via parathyroid hormone-related protein (PTHrP) (Amano et al., 2009; Nishimura, Hata, Ono, et al., 2012). However, PTHrP was not strongly regulated in our data set. Furthermore, our RT-qPCR revealed that only V602I significantly upregulated *SOX9*, and the RNAseq data showed that *SOX9* had a smaller fold change compared to other chondrogenic genes, such as *GDF5, COL6A1, COL6A3,* and *COMP*. In fact, these genes, which were upregulated in V620I- and T89I-hiPSC-derived chondrocytes, have a pro-chondrogenic but anti-hypertrophic phenotype (Caron et al., 2020; Chu et al., 2017; Hecht & Sage, 2006). Therefore, these results suggest additional and alternative pathways to *FST* and *SOX9* that are responsible for the V620I and T89I skeletal dysplasias.

Our results are also generally consistent with previous reports on the effects of other TRPV4 mutations such as lethal and non-lethal metatropic dysplasia-causing I604M (Saitta et al., 2014) and L619F (Nonaka et al., 2019). The data also reveal potential differences in the effects of these varying TRPV4 mutations on cell electrophysiology or differentiation. For example, we saw an increase in *SOX9* expression in V620I, while no change in T89I. Gain-of-function mutation L619F also increased *SOX9* expression (Nonaka et al., 2019), while I604M, which has been reported to not alter conductivity like T89I (Sun, 2012), decreased *SOX9* (Saitta et al., 2014). I604M also decreased *COL2A1*, *COL10A1*, and *RUNX2* expression consistent with our T89I results (Saitta et al., 2014). Intriguingly, the L619F mutation was reported to increase Ca^2+^ signaling with activation via a TRPV4 agonist (Nonaka et al., 2019). However, we observed that V620I and T89I had significantly reduced Ca^2+^ signaling compared to WT in response to chemical agonist GSK101, as confirmed by both confocal imaging and patch clamping. These results highlight that TRPV4 mutations have heterogeneous effects on downstream signaling pathways and thus lead to diverse disease phenotypes, despite similar classification of these mutations as “gain-of-function.” It is also important to note that in previous studies, chondrogenic differentiation of iPSCs (Saitta et al., 2014) or dental pulp cells (Nonaka et al., 2019) were performed in short-term micromass culture, and not long-term pellet culture as in our study, potentially leading to different levels of chondrogenesis and maturation of the cells.

Our transcriptomic analysis showed significant changes in various *HOX* family genes due to TRPV4 mutations, suggesting a potential role of these genes in maintaining the immature, chondrogenic phenotype in the mutated lines. At both day 28 and 56, the top 25 up-regulated genes in the V620I and T89I lines included genes from the anterior *HOX* family (Iimura & Pourquié, 2007; Seifert, 2015). The high expression of anterior *HOX* genes indicates that the mutants are maintaining the chondrocytes in an early developmental stage with axial patterning. At day 28 and 56, *HOXA2*, *HOXA3*, and *HOXA4* were in the top upregulated genes, with *HOXA4* having the largest fold change. Interestingly, gain-of-function mutations or overexpression of *HOXA2, HOXA3*, and *HOXA4* impair chondrogenesis, limit skeletal development, decrease endochondral ossification regulators, and delay mineralization in animal models (Creuzet, Couly, Vincent, & Le Douarin, 2002; Deprez, Nichane, Lengele, Rezsohazy, & Nyssen-Behets, 2013; Kanzler, Kuschert, Liu, & Mallo, 1998; Li, 2006; Massip et al., 2007; Seifert, 2015). *HOXA5* was also highly upregulated at both day 28 and 56, and mutations in this gene showed disordered patterning of limb bud development (Pineault & Wellik, 2014). Finally, the rib and spine phenotypes associated with brachyolmia and metatropic dysplasia could be contributed to the altered expression of *HOXA4* to *HOXA7* as it has been shown that these genes are associated with rib and spine patterning, and alterations in expression have led to defects (F. Chen, Greer, & Capecchi, 1998; Wellik, 2009). The only up-regulated posterior *HOX* genes were *HOXC8* and *HOXD8* at day 56 (Iimura & Pourquié, 2007; Seifert, 2015). The absence of posterior *HOX9, HOX11,* and *HOX13*, which are associated with limb development and hypertrophic *RUNX2/3* expression (Pineault & Wellik, 2014; Qu, Palte, Gontarz, Zhang, & Guilak, 2020), may be at least partially responsible for the improper development in skeletal dysplasias. Interestingly, many links have been identified between *HOX* genes and TGFβ3-family signaling, specifically through SMAD proteins, both within skeletal development and other processes (e.g., murine lung development) (Li, 2006; Li & Cao, 2003; Volpe et al., 2013).

In fact, TRPV4 and TGF-β signaling have recently been shown to interact, with effects specific to the order in which they occur (Nims et al., 2021; O’Conor et al., 2014; Woods et al., 2021). Consistent with previous finding with hiPSCs housing the I604M TRPV4 mutations (Saitta et al., 2014), the altered TRPV4 activity in our hiPSC-derived chondrocytes could be altering their response to the TGFβ3 and BMP4 treatments. Furthermore, the V620I and T89I mutations increased expression of *HTRA1,* which has been shown to bind to and alter the response to members of the TGFβ family (Polur, Lee, Servais, Xu, & Li, 2010). Furthermore, *TGFβ3* and *TWIST,* which is downstream of TGFβ3-signaling, were both upregulated in TRPV4- mutated hiPSC-derived chondrocytes. It has been reported that *TGFB3* expression and signaling prevent osteoblastogenesis of mesenchymal stem cells (Nishimura, Hata, Matsubara, et al., 2012; Nishimura, Hata, Ono, et al., 2012), while *TWIST* inhibits hypertrophy regulators *RUNX2* and *FGFR2* (Michigami, 2014; Miraoui & Marie, 2010). Therefore, another mechanism of hypertrophic dysregulation with these mutations could be altered response to TGFβ family signaling.

Furthermore, significantly lower expression of *ALPL, AMELX, IFITM5, GSTA1, IBSP*, and *MEPE* in mutated chondrocytes compared to WT suggest that mutated cells had altered response to BMP4-induced hypertrophy. Indeed, mutations in *ALPL* have been shown to lead to hypophosphatasia with deformed long bones (Taillandier et al., 2015), while mutations in *IFITM5* lead to osteogenic imperfecta (Hanagata, 2016). Our results indicate a connection between these genes and delayed endochondral ossification in chondrocytes bearing V620I and T89I mutations; however, how the expression levels of these genes are associated with TRPV4 function and mutations still warrants further investigation.

Another gene increased in BMP4-treated WT, but less in mutants, was *GSTA1,* which produces the antioxidant glutathione (C.-T. Chen, Shih, Kuo, Lee, & Wei, 2008; Hayes, Flanagan, & Jowsey, 2005). The TRPV4-mutated chondrocytes also had significant downregulation of catalase (*CAT*), another antioxidizing gene (C.-T. Chen et al., 2008). Interestingly, BMP4 treatment of T89I-mutated chondrocytes was able to significantly increase *CAT* expression, potentially indicating an association between antioxidant expression and maturation. One study observed that chondrocyte maturation is associated with decreasing catalase (Morita et al., 2007). However, many others report that reactive oxygen species (ROS; e.g., H_2_O_2_), which can be removed by *CAT* and *GSTA1*, prevent endochondral ossification, potentially via inhibition of the hedgehog pathways (Atashi, Modarressi, & Pepper, 2015; C.-T. Chen et al., 2008; Fragonas et al., 1998). Interestingly, *IHH* also had the lowest expression level in our T89I mutant chondrocytes. These findings suggest that decreased expression of *CAT* and *GSTA1* in TRPV4 mutants may also be involved in dysregulating endochondral ossification in these cells.

The decrease in mechanical properties with increased basal current of the V620I mutant was unexpected since TRPV4 activation was previously shown to increase matrix production and properties (O’Conor et al., 2014). Furthermore, genes uniquely upregulated in V620I were associated with interferon type I (IFNβ). IFNβ has been reported to decrease inflammatory markers and matrix degradation (Hu, Ho, Lou, Hidaka, & Ivashkiv, 2005; Palmer et al., 2004; van Holten et al., 2004; Zhao et al., 2014), despite the decrease in moduli observed in the day-42 V620I chondrogenic pellets. Interestingly, a study comparing bone marrow-derived MSCs from healthy and systemic lupus erythematous patients found that IFNβ inhibited osteogenesis via suppression of *RUNX2* and other osteogenic genes (Gao, Liesveld, Anolik, McDavid, & Looney, 2020). Highlighting a potential, unique regulator of the delayed hypertrophy in V620I leading to brachyolmia. In contrast, T89I became more unique at later time points of chondrogenesis and was not associated with many of the same biological processes as WT and V620I, especially those regarding endochondral ossification, when treated with BMP4. This, in conjunction with the high number of unique DEGs, represents a potential inhibition of hypertrophy, particularly in response to BMP4 treatment, with the T89I mutation leading to severe metatropic dysplasia.

Here, we present multiple putative genes and pathways that could be involved in delaying, and potentially inhibiting, chondrocyte hypertrophy in V620I- and T89I-TRPV4 mutants. It should be noted, however, that this study has some potential limitations. It is well-recognized that Wnt/β-catenin signaling plays an important role in chondrocyte hypertrophy (Hou et al., 2019; Huang, Zhong, Hendriks, Post, & Karperien, 2018; Michigami, 2014). However, we may be preventing some hypertrophy since our chondrogenic protocol uses a pan-Wnt inhibitor to prevent off-target differentiation and promote a homogenous chondrocyte population (Wu et al., 2021). Nevertheless, our WT chondrocytes, but not TRPV4 mutants, exhibited hypertrophic differentiation with BMP4 treatment, suggesting that DEGs/pathways detected in our sequencing analysis are still robust. Since this study focuses on TRPV4 gain-of-function mutations, future studies could fully or partially inhibit TRPV4 signaling to determine if that would increase similarity between the mutant and WT lines at various stages of chondrogenic and hypertrophic differentiation. Additionally, this study only activated TRPV4 using the pharmacological activator GSK101. Other future experiments could activate the channel osmotically or with mechanical loading to investigate additional differences in TRPV4 function leading to skeletal dysplasias during development.

In summary, our study found that dysregulated skeletal development in the V620I- and T89I-TRPV4 dysplasias is likely due, at least in part, to delayed and inhibited chondrocyte hypertrophy. The gain-of-function mutations may lead to increased *HOX* gene expression, altered TGFβ signaling, decreased hypertrophic and biomineralization gene expression (e.g., *ALPL, AMELX, IFITM5, IBSP*, and *MEPE*), and genes regulating hedgehog pathways and ROS accumulation (e.g., *GSTA1, CAT*). These findings lay a foundation for the development of therapeutics for these diseases and give insights into the regulation of endochondral ossification via TRPV4.

## MATERIALS AND METHODS

### hiPSC culture

The BJFF.6 (BJFF) human iPSC line (Washington University Genome Engineering and iPSC Center (GEiC), St. Louis, MO), was used in this study as the isogenic-wildtype control. CRISPR-Cas9 gene editing was used to create the V620I and T89I mutations in the BJFF cell line as described previously (S. S. Adkar et al., 2019). The hiPSCs were maintained on vitronectin (VTN-N; cat. num. A14700; Thermo Fisher Scientific, Waltham, MA)-coated plates in Essential 8 Flex medium (E8; cat. num. A2858501; Gibco, Thermo Fisher Scientific, Waltham, MA). Medium was changed daily until cells were passaged at 80-90% confluency (medium supplemented with Y-27632 [cat. num. 72304; STEMCELL Technologies, Vancouver, Canada] for 24 hours) or induced into mesodermal differentiation at 30-40% confluency.

### Mesodermal differentiation

The hiPSCs were differentiated through the mesodermal pathway as previously described (S. S. Adkar et al., 2019; Dicks et al., 2020; Wu et al., 2021). In brief, cells were fed daily with different cocktails of growth factors and small molecules for twelve days in mesodermal differentiation medium and driven through the anterior primitive streak (1 day; 30 ng/ml Activin [cat. num. 338-AC; R&D Systems, Minneapolis, MN], 20 ng/ml FGF2 [cat. num. 233-FB-025/CF; R&D Systems, Minneapolis, MN], 4 µM CHIR99021 [cat. num. 04-0004-02; Reprocell, Beltsville, MD]), paraxial mesoderm (1 day; 20 ng/ml FGF2, 3 µM CHIR99021, 2 µM SB505124 [cat. num. 3263; Tocris Bioscience, Bristol, UK], 4 µM dorsomorphin [DM; cat. num. 04-0024; Reprocell, Beltsville, MD]), early somite (1 day; 2 µM SB505124, 4 µM dorsomorphin, 500 nM PD173074 [cat. num. 3044; Tocris Bioscience, Bristol, UK], 1 µM Wnt-C59 [cat. num. C7641-2s; Cellagen Technologies, San Diego, CA]), and sclerotome (3 days; 1 µM Wnt-C59, 2 µM purmorphamine [cat. num. 04-0009; Reprocell, Beltsville, MD]) into chondroprogenitor cells (6 days; 20 ng/ml BMP4 [cat. num. 314-BP-010CF; R&D Systems, Minneapolis, MN]). Mesodermal differentiation medium had a base of Iscove’s Modified Dulbecco’s Medium, glutaMAX (IMDM; cat. num. 31980097; Gibco, Thermo Fisher Scientific, Waltham, MA) and Ham’s F-12 nutrient mix, glutaMAX (F12; cat. num. 31765092; Gibco, Thermo Fisher Scientific, Waltham, MA) in equal parts supplemented with 1% penicillin-streptomycin (P/S; cat. num. 15140122; Gibco, Thermo Fisher Scientific, Waltham, MA), 1% Insulin-Transferrin-Selenium (ITS+; cat. num. 41400045; Gibco, Thermo Fisher Scientific, Waltham, MA), 1% chemically defined concentrated lipids (cat. num. 11905031; Thermo Fisher Scientific, Waltham, MA), and 450 µM 1-thioglycerol (cat. num. M6145; Millipore Sigma, St. Louis, MO). The chondroprogenitor cells were then disassociated for chondrogenic differentiation.

### Chondrogenic differentiation with 3D pellet culture

Cells were differentiated into chondrocytes using a high-density, suspension pellet culture (S. S. Adkar et al., 2019; Dicks et al., 2020; Wu et al., 2021). In summary, cells were resuspended in chondrogenic medium: Dulbecco’s Modified Eagle Medium/F12, glutaMAX (DMEM/F12; cat. num. 10565042; Gibco, Thermo Fisher Scientific, Waltham, MA), 1% P/S, 1% ITS+, 1% Modified Eagle Medium (MEM) with nonessential amino acids (NEAA; cat. num. 11140050; Gibco, Thermo Fisher Scientific, Waltham, MA), 0.1% dexamethasone (Dex; cat. num. D4902; Millipore Sigma, St. Louis, MO), and 0.1% 2-Mercaptoethnol (2-ME; cat. num. 21985023; Gibco, Thermo Fisher Scientific, Waltham, MA) supplemented with 0.1% L-ascorbic acid (ascorbate; cat. num. A8960; Millipore Sigma, St. Louis, MO), 0.1% L-proline (proline; cat. num. P5607; Millipore Sigma, St. Louis, MO), 10 ng/ml human transforming growth factor-β3 (TGFβ3; cat. num. 243-B3-010/CF; R&D Systems, Minneapolis, MN), 1 µM Wnt-C59, and 1 µM ML329 (cat. num. 22481; Cayman Chemical, Ann Arbor, MI) at 5 x 10^5^ cells/mL. One mL of the cell solution was added to a 15 mL-conical tube (cat. num. 430790; Corning, Corning, NY) and centrifuged to form the spherical pellets. Pellets were fed every 3-4 days with complete chondrogenic medium until the desired time point. Several timepoints of the chondrogenic pellets were used to study chondrocyte maturation (7, 14, 28, and 42 days), mechanical properties (28 and 42 days), hypertrophy (28 days) or, after digestion to single cell day-28 chondrocytes, on Ca^2+^ signaling in response to pharmacological activation of TRPV4.

### BMP4 treatment to promote hypertrophic differentiation

Some day-28 pellets were also further differentiated for an additional 4 weeks to examine the effects of the mutations on chondrocyte hypertrophy. Pellets were cultured with complete chondrogenic medium with either TGFβ3 (10 ng/mL) alone, BMP4 (50 ng/mL) alone, or a combination of TGFβ3 (10 ng/mL) and BMP4 (50 ng/mL).

### Dissociation of chondrogenic pellets to obtain single cell hiPSC-derived chondrocytes

To isolated hiPSC-derived chondrocytes, day-28 chondrogenic pellets were rinsed and placed in an equal volume (1 pellet per 1 mL) of digestion medium (0.4% w/v type II collagenase [cat. num. LS00417; Worthington Biochemical, Lakewood, NJ] in DMEM/F12 with 10% fetal bovine serum [FBS; cat. num. S11550; Atlanta Biologicals, R&D Systems, Minneapolis, MN]). The tubes were placed on an orbital shaker at 37°C and vortexed every 20 minutes for approximately 2 hours. Once the tissue was digested and could no longer be seen by the naked eye, the digestion medium was neutralized in DMEM/F12 medium containing 10% FBS. These cells were used for patch clamping and confocal experiments.

### TRPV4 agonists and antagonists

Solutions were prepared immediately before experiments and held at room temperature. GSK1016790A (GSK101; cat. num. G0798; Sigma Aldrich, St. Louis, MO) and/or GSK205 (cat. num. AOB1612 1263130-79-5; AOBIOUS, Gloucester, MA), in addition to DMSO for a vehicle control, were added to assay buffer (Hanks’ Balanced Salt Solution [HBSS; cat. num. 14025076; Gibco, Thermo Fisher Scientific, Waltham, MA] with 2% HEPES [cat. num. 15630130; Gibco, Thermo Fisher Scientific, Waltham, MA]) at 2x the desired concentration (20 nM GSK101, 40 µM GSK205). Solutions were made at 2x the desired concentration because they would be mixed at an equal volume of assay buffer after capturing a baseline fluorescence in Ca^2+^ signaling experiments.

### Patch clamping

Isolated chondrocytes were kept on ice and used for patching within 36 hours. Patch-clamp experiments were carried out at RT under two conditions. Single-channel measurements were made in excised inside-out membrane patches in a symmetric potassium chloride (KCl) solution (148mM KCl, 1mM K_2_EDTA, 1mM EGTA, 10mM HEPES, pH 7.4). Channel activation was achieved by bath perfusion with the same buffer solution containing 10 nM GSK101. Blocking was performed using the same buffer solution supplied with both 10 nM GSK101 and 20 µM GSK205. Recordings were made at -30mV membrane. Whole-cell currents were recorded using an external sodium chloride (NaCl) solution (150 mM NaCl, 5 mM KCl, 1 mM EGTA, 10 mM Glucose, 10 mM HEPES, and 10 µM free Ca^2+^) and KCl pipette solution as used for single-channel recordings. Inhibition of basal currents was performed by pre-incubation of the cells in external solution supplied with 20 µM GSK205 for 20 min before patching; the drug was also present in the bath at the same concentration during the experiment. Data were acquired at 3 kHz, low-pass filtered at 1 kHz with Axopatch 1D patch-clamp amplifier and digitized with Digidata 1320 digitizer (Molecular Devices, San Jose, CA). Data analysis was performed using the pClamp software suite (Molecular Devices, San Jose, CA). Pipettes with 2.0-4.0 MOhm resistance in symmetric 150 mM KCl buffer were pulled from Kimble Chase 2502 soda lime glass with a Sutter P-86 puller (Sutter Instruments, Novato, CA).

### Confocal imaging of Ca^2+^ signaling

hiPSC-derived chondrocytes from digested pellets were plated in DMEM medium containing 10% FBS at 2.1 x 10^4^ cells/cm^2^ in 35 mm-dishes for 6-8 hours to allow the cells to adhere without dedifferentiating. Cells were then rinsed and stained for 30 min with Fluo-4 AM (cat. num. F14201; Thermo Fisher Scientific, Waltham, MA), Fura Red AM ((cat. num. F3021; Thermo Fisher Scientific, Waltham, MA), and sulfinpyrazone (cat. num. S9509-5G; Sigma Aldrich, St. Louis, MO) with 20 mM GSK205 or 1000x DMSO (vehicle control). The dye solution was replaced with assay buffer before imaging cells on a confocal microscope (LSM 880; Zeiss, Oberkochen, Germany) at baseline for the first 100 frames (approximately 6 min). Then, an equal volume of a 2x solution of GSK101 or GSK101 and GSK205 was added, and imaging continued for an additional 300 frames (approximately 20 min). Fiji software (ImageJ, version 2.1.0) was used to locate cells and quantify the ratiometric fluorescence intensity (Intensity_fluo-4_/Intensity_fura red_). In brief, .czi files were imported into Fiji and the channels were split. After applying the median filter, the image calculator divided the green channel by the red. A Z-projection was performed based on the maximum fluorescence of the red channel (to ensure that all cells were identified even in groups were there was no increase in Ca^2+^ signaling). A threshold and watershed binary were then applied, and measurements were set for a cell size of 100-infinity. Outlines were projected, and the mean fluorescence of each cell was measured over time. The average fluorescence was plotted for all the cells in the group over time. Area under the curve and time of response were calculated to quantify differences between groups. Cells were classified as responders if they had a fluorescence greater than the baseline mean plus 3 times the standard deviation in at least a quarter of the frames. Time of response was the time of the first frame in which the cell responded for at least 2 consecutive frames. The fluorescence was measured for all the cells in the frame of view as technical replicates for 2 experimental replicates.

### AFM measurement of neocartilage mechanical properties

Day-28 and day-42 hiPSC-derived pellets were rinsed in PBS and snap frozen in optimal cutting temperature (OCT; cat. num. 4583; Sakura Finetek, Torrance, CA) medium and stored at -80 °C. Pellets were cryosectioned using cryofilm (type 2C(10); Section-Lab, Hiroshima, Japan) in multiple different regions of the pellet (i.e., zones). The 10 µm cryosection with cryofilm was fixed on a microscope slide using chitosan and stored at 4 °C overnight. The next day, cryosections were mechanically loaded using an atomic force microscopy (AFM, MFP-3D Bio, Asylum Research, Goleta, CA) as previously described (Votava, Schwartz, Harasymowicz, Wu, & Guilak, 2019). Briefly, the samples were tested in PBS at 37 °C to maintain hydration and mimic physiologic conditions, respectively. The sections were mechanically probed using a silicon cantilever with a spherical tip (5μm diameter, k∼7.83 N/m, Novascan Technologies, Ames, IA). An area of 10 μm^2^ with 0.5 μm intervals (400 indentations) was loaded to 300 nN with the loading rate of 10 μm/sec. Multiple locations from different sites of each zone and pellet were loaded as replicates. The curves obtained from AFM were imported into a custom written MATLAB code to determine the mechanical properties of the pellets. Using contact point extrapolation, the contact point between the cantilever’s tip and the tissue was detected, and the elastic modulus was calculated using a modified Hertz model (Darling, Wilusz, Bolognesi, Zauscher, & Guilak, 2010; Darling, Zauscher, & Guilak, 2006; Votava et al., 2019; Wilusz, Zauscher, & Guilak, 2013; Zelenski et al., 2015).

### Histology

Chondrogenic pellets at days 7, 14, 28, 42, and 56 (with and without BMP4) were fixed and dehydrated in sequential steps of increasing ethanol and xylene solutions until embedded in paraffin wax. Wax blocks were cut into 8 µm sections on microscope slides for histological and immunohistochemical analysis. Slides were rehydrated in ethanol and water and the nuclei were stained with Harris hematoxylin and sGAGs with Safranin-O. Antigen retrieval was performed on rehydrated slides followed by blocking, the addition of primary and secondary antibodies, and AEC development to label collagen proteins (COL1A1, COL2A1, COL6A1, and COL10A1) and Vector Hematoxylin QS counterstain.

### Biochemical analysis

Chondrogenic pellets at days 7, 14, 28, and 42 were washed with PBS and digested in papain overnight at 65°C. sGAG and dsDNA content were measured using the dimethylmethylene blue (DMMB) and PicoGreen assays (Quant-iT™ PicoGreen™ dsDNA Assay Kit; cat. num. P7589; Thermo Fisher Scientific, Waltham, MA) respectively. sGAG content was normalized to dsDNA. Three to four independent experiments were performed with 3-4 technical replicates per group.

### Western blot

Day-56 pellets treated with TGFβ3, TGFβ3+BMP4, or BMP4 were digested to single cells, as described above, and lysed in RIPA buffer (cat. num. 9806S; Cell Signaling Technology, Danvers, MA) with protease inhibitor (cat. num. 87786; Thermo Fisher Scientific, Waltham, MA). Protein concentration was then measured using the BCA Assay (Pierce). Twenty micrograms of proteins for each well were separated on 10% sodium dodecyl sulfate-polyacrylamide gel electrophoresis gel with pre-stained molecular weight markers (cat. num. 161-0374; Bio-Rad, Hercules, CA) and transferred to a polyvinylidene fluoride (PVDF) membrane. The PVDF membrane blot was cut through the line at 50 kD. Two blots were incubated overnight at 4 °C with the primary antibodies: anti-COL10A1 (1:500; cat. num. PA5-97603; Thermo Fisher Scientific, Waltham, MA) and anti-GAPDH (1:30000; cat. num. 60004-1-Ig; Proteintech, Rosemont, IL), as the loading control. TidyBlot-Reagent-HRP (1:1000; cat. num. 147711; Bio-RAD, Hercules, CA) and horse anti-mouse IgG secondary antibody (1:3000; cat. num. 7076; Cell Signaling, Danvers, MA) were then used respectively. Immunoblots were imaged and analyzed using the iBright FL1000 Imaging System (Thermo Fisher Scientific, Waltham, MA).

### RNA isolation

Chondrogenic pellets at days 7, 14, 28, 42, and 56 were washed with PBS, lysed, snap frozen, and homogenized. RNA was isolated using the Total RNA Purification Plus Kit (cat. num. 48400; Norgen Biotek, Thorold, Canada) and used immediately for either RT-qPCR or RNA-seq.

### Gene expression with RT-qPCR

Isolated RNA was reverse transcribed into cDNA. The cDNA was used to run real-time, quantitative PCR using Fast SYBR green. Gene expression was analyzed using the ΔΔC_T_ method with hiPSC as the reference time point and *TBP* as the housekeeping gene (Livak & Schmittgen, 2001). Three to four independent experiments were performed with 3-4 technical replicates per group. Primers can be found in the Fig. 3 – Supplemental Table 1.

### Genome-wide mRNA sequencing

Isolated RNA was treated with DNase (cat. num. 25720; Norgen Biotek, Thorold, Canada) and cleaned (cat. num. 43200; Norgen Biotek, Thorold, Canada) according to manufacturer instructions prior to submitting to the Genome Technology Access Center at Washington University in St. Louis (GTAC). Libraries were prepared according to manufacturer’s protocol. Samples were indexed, pooled, and sequenced at a depth of 30 million reads per sample on an Illumina NovaSeq 6000. Basecalls and demultiplexing were performed with Illumina’s bcl2fastq software and a custom python demultiplexing program with a maximum of one mismatch in the indexing read. RNA-seq reads were then aligned to the Ensembl release 76 primary assembly with STAR version 2.5.1a (Dobin et al., 2013). Gene counts were derived from the number of uniquely aligned unambiguous reads by Subread:featureCount version 1.4.6-p5 (Liao, Smyth, & Shi, 2014). Isoform expression of known Ensembl transcripts were estimated with Salmon version 0.8.2 (Patro, Duggal, Love, Irizarry, & Kingsford, 2017). Sequencing performance was assessed for the total number of aligned reads, total number of uniquely aligned reads, and features detected. The ribosomal fraction, known junction saturation, and read distribution over known gene models were quantified with RSeQC version 2.6.2 (Wang, Wang, & Li, 2012).

### Transcriptomic analysis of sequencing datasets

R and the DESeq2 package were used to read un-normalized gene counts, and genes were removed if they had counts lower than 200 (Love, Huber, & Anders, 2014). Regularized-logarithm transformed data of the samples were visualized with the *Pheatmap* package (Kolde, 2015) function on the calculated Euclidean distances between samples or with the *ggplot2* package (Wickham, 2009) to create a principle component analysis (PCA). The transformed data was also used to determine the top 5000 most variable genes across the samples. The replicates, from DESeq data, for each group were averaged together, and the up- and down-regulated differentially expressed genes (DEGs) were determined. The total number of DEGs was plotted using GraphPad Prism. At day 28, the V620I and T89I lines were compared to WT. At day 56, TGFβ3-treated V620I and T89I were compared to TGFβ3-treated WT, and BMP4-treated groups were compared to their respective TGFβ3-treated group of the same line (e.g., BMP4-treated WT vs. TGFβ3-treated WT). Genes were considered differentially expressed if adjusted p value (p_adj_) < 0.1 and log_2_(fold change) ≥ 1 or ≤ -1. The intersecting and unique DEGs were determined and plotted with the *intersect* and *setdiff,* and *venn.diagram* (*VennDiagram* package (H. Chen & Boutros, 2011)) functions. The fold changes of common chondrogenic, hypertrophic, growth factor, Ca^2+^ signaling, and off-target genes, in the top 5000 most variable genes, were plotted using the *pheatmap* function. The top 25 most up-regulated and down-regulated for each group, based on log_2_(fold change), and the log_2_(fold change) of that gene for the other group(s) were also plotted with the *pheatmap*. Gene lists (e.g., intersected genes, genes upregulated with BMP4 treatment) were entered into g:profiler to determine associated Gene Ontology (GO) Biological Processes, Molecular Functions, Cellular Components, KEGG pathways, Reactome pathways, and Human Phenotype (HP) Ontologies (Raudvere et al., 2019). The negative log_10_ of the adjusted p value for each term was plotted with GraphPad Prism or using a function to scale circle diameter to the p value in Illustrator.

The gap statistic method determined the ideal number of clusters resulting from BMP4 treatment was either 1 or 9. We then performed k-means clustering with 9 clusters and plotted the gene expression trends for each gene within the cluster with the average expression trend overlaying for each cell line of the largest cluster using the *tidyverse* package (Altman & Krzywinski, 2017). The genes in each cluster, with the normalized counts for each group, are listed in Supplemental File 1. The largest cluster was plotted using the Cytoscape String app’s protein interaction to create a protein-protein network (Doncheva, Morris, Gorodkin, & Jensen, 2019; Shannon, 2003). Using the average log fold change with BMP4 treatment across lines, the network was propagated using the Diffusion app, and functional enrichment with EnrichmentMap was performed on the network (Merico, Isserlin, Stueker, Emili, & Bader, 2010). We then created a network connecting the genes to their associated genes with black lines and to their associated Gene Ontology processes using grey lines. We colored the gene circles with three colors representing the log fold change of that gene in each line. The white arrows were added to the color scale legend to indicate maximum log fold change for each line.

### Statistical analysis

Data were graphed and analyzed using GraphPad Prism (Version 9.1.0). Outliers were removed from the data using the ROUT method (Q = 1%), and the data were tested for normality with the Shapiro-Wilk test (α = 0.05). For RT-qPCR, normally distributed data were analyzed within each time point using a Brown-Forsythe and Welch one-way ANOVA with multiple comparisons (mean of each column, cell line, with every other column). A Kruskal-Wallis test was used if data was not normally distributed. For biochemical analysis, mechanical properties, and area under the curve, and time of response, data were analyzed using an ordinary two-way ANOVA, comparing each cell with all other cells, with Tukey’s post-hoc test. Area under the curve was quantified for plots over time considering a baseline of Y=0, ignoring peaks less than 10% of the distance from minimum to maximum Y, and all peaks going over the baseline.

## ACKNOWLEDGMENTS

This work was supported by Shriners Hospitals for Children – St. Louis, the National Institute of Health (R01 AG46927, R01 AG15768, R01 AR072999, R00 AR075899, P30 AR073752, P30 AR074992, T32 DK108742, T32 EB018266, and CTSA grant UL1 TR002345). We would like to thank the Washington University Genome Engineering and iPSC Center and the Genome Technology Access Center (GTAC) for their assistance with the CRISPR-Cas9 editing and RNA sequencing, respectively. We would also like to thank Dr. Monica Sala-Rabanal for assistance and advice in the initial aspects of this study.

